# Imipramine binds to Amyloid-β (1-42) monomers in vitro, as shown by NMR spectroscopy

**DOI:** 10.64898/2026.08.24.746779

**Authors:** James Beham, Noah R. Johnson, Beat Vögeli, Morkos A. Henen, Liliya Vugmeyster

## Abstract

Imipramine is known as an older generation tricyclic antidepressant drug. It has been identified in prior studies that imipramine blocks Apolipoprotein E4 (ApoE4)-induced amyloid-β(Aβ) aggregation and is associated with an improved AD diagnosis [Johnson *et al. Alzheimer’s Research & Therapy, 2022*, 14:88]. Using NMR methods such as ^1^H-^1^H NOESY and Saturation Transfer Difference Spectroscopy, we demonstrate the binding of Aβ monomers to imipramine when the full-length Aβ (1-42) sequence is considered. The more abundant but less toxic form, Aβ (1-40) does not show interaction with imipramine.

## Introduction

Imipramine is known as an older generation tricyclic antidepressant drug, with structure shown in Fig. S1. It has been identified in prior studies that imipramine blocks Apolipoprotein E4 (ApoE4)-induced amyloid-β(Aβ) aggregation and is associated with an improved AD diagnosis.^1^ Out of several FDA-approved drug candidates, it was shown to reduce neurodegeneration and improve cognition in individuals carrying the ApoE genetic mutation.^1^

The goal of this work is to directly test its binding to amyloid-β monomers in vitro. We will probe both the full length 42 residues sequence (Aβ42) which is considered most toxic, and the shorter truncated version Aβ40, which is most abundant.^2,3^ We apply well-established approaches such as two-dimensional 1H-1H NOESY and Saturation Transfer Difference (STD) Spectroscopy.^4^ Both experiments rely on the Nuclear Overhauser Effect. In a classic 2D NOESY spectrum, binding between the ligand and protein can be detected by a change of sign in the cross-peaks of the ligand. A free ligand behaves as a small molecule in the fast-tumbling regime, which results in a difference in the phase between diagonal and cross peaks (negative cross-peaks which are of opposite sign to the diagonal). On the other hand, a ligand bound to a macromolecule such as Aβ, behaves as a large molecule due to the slower tumbling and hence cross and diagonal peaks for this ligand will have the same phase (positive cross-peaks, with the same sign as the diagonal).^5,6^ In the STD experiment, low power irradiation is applied to the region of the spectrum containing protein signal, which spreads through the entire macromolecule due to spin-diffusion and results in the magnetization transfer to the ligand, if it is transiently bound to the macromolecule.^4^

## Results

### Imipramine binds Aβ monomer

As stated in the Introduction, if the ligand is bound to a macromolecule, it behaves as a large molecule, as reflected in the same sign of the diagonal and cross-peaks in the 2D NOESY spectrum. This is indeed the case for free imipramine (Fig. 1a, negative cross-peaks are in red, positive peaks are in blue) and imipramine bound to Aβ42 monomer (Fig. 1b). Additionally, there is a clear broadening of the cross-peaks in the presence of Aβ42 (Fig. S2), serving as additional confirmation of binding. We then proceeded with the STD method, based on the transfer of saturation of resonances from the macromolecule to the ligand. The experiment was performed twice, where the gaussian saturation pulse was applied on resonance to target the Aβ42 monomer protein methyl resonances and then off-resonance as a control. The two experiments were subtracted, and the STD effect was observed. The spectrum of free imipramine is shown in Fig. 2. where we observe zero STD signal at concentrations below 400 μM, as expected for a negative control. In the presence of Aβ42 monomer, the binding leads to non-zero STD signal of imipramine indicating binding and distance proximity (Fig. 2). To confirm this binding, we performed the STD measurements at three different temperatures (4, 10, and 20 °C) and we observed non-zero STD signal in all cases (Fig. S3). Using N^15^-labeled Aβ42, we also collected heteronuclear single-quantum correlation (HSQC) NMR spectra of Aβ42 monomers alone (Fig. 3) and those in the presence of imipramine. For a structured protein, changes in the backbone resonance positions are expected upon ligand binding for the sites affected by the ligand. In our case we did not observe any chemical shift changes. As Aβ42 is disordered, the subtle changes in the conformational ensemble are not visible with this technique and suggest that binding might also be more pronounced at the side-chain sites. It is interesting to note that in contrast to the case of Aβ42, the imipramine NOESY spectrum in the presence Aβ40 monomers (Fig. S4) does not display the features characteristic of binding: the cross-peaks display neither the change of sign nor the broadening. Thus, the binding is restricted to Aβ42.

**Figure 1.**
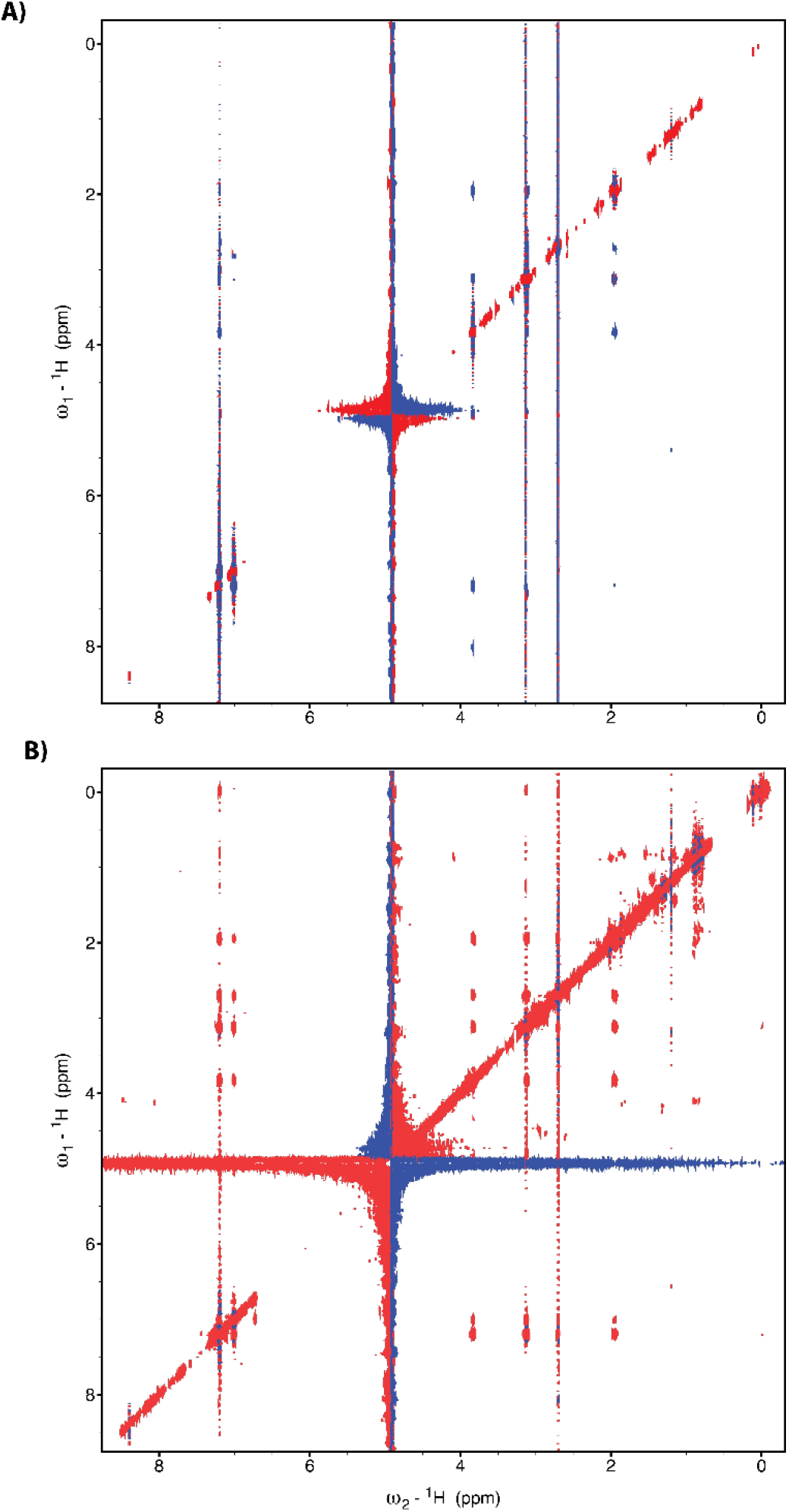
^1^H-^1^H NOE spectra of A) imipramine at 200 μM concentration and B) imipramine at 200 μM concentration in the presence of 25 μM Aβ42 monomers, at 10 oC. The mixing time was 300 ms. The red and blue colors indicated negative and positive intensities, respectively. The binding can be seen by the change of sign of the cross-peak of imipramine upon binding, as well as the widths of the peaks.

**Figure 2.**
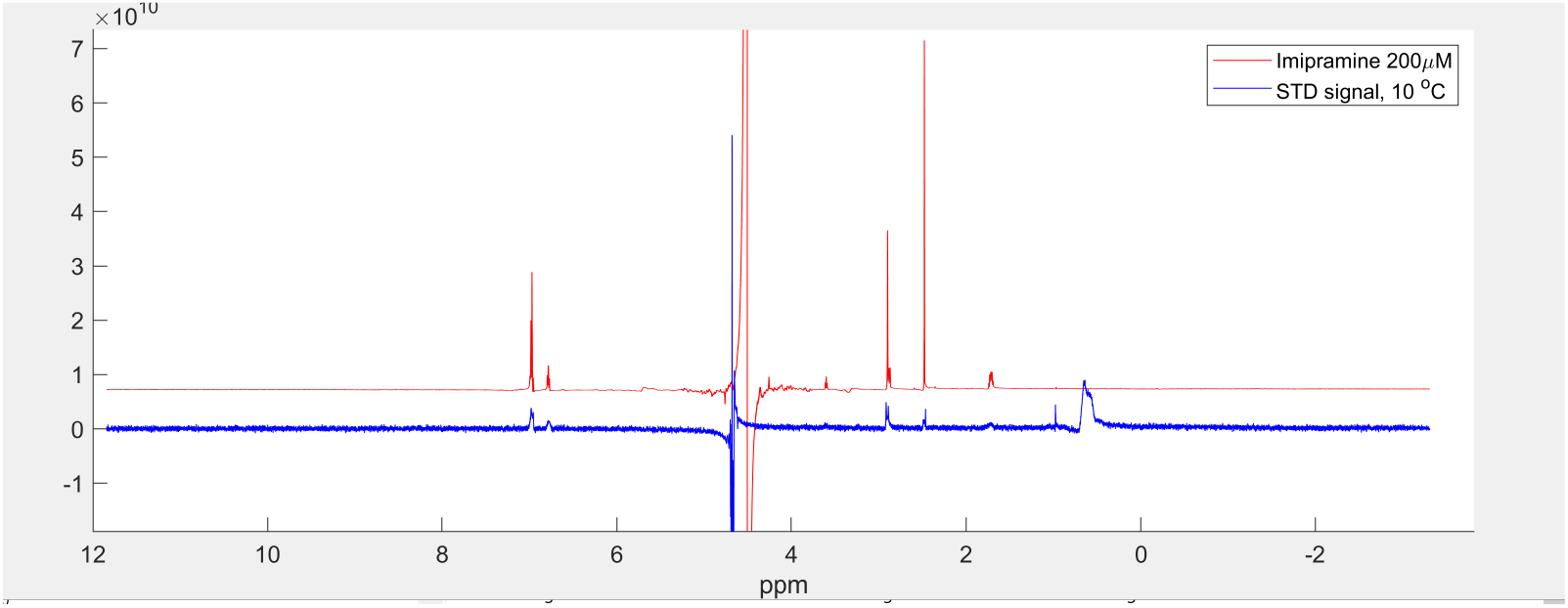
^1^H NMR saturation transfer difference data for Aβ1-42. Concentrations were 20 μM Aβ42 and 200 μM imipramine, at 10 oC. Intensity in arbitrary units versus chemical shift. Red line: imipramine in isolation, blue line: STD signal, scaled by a factor of 23 relative to the red trace.

**Figure 3.**
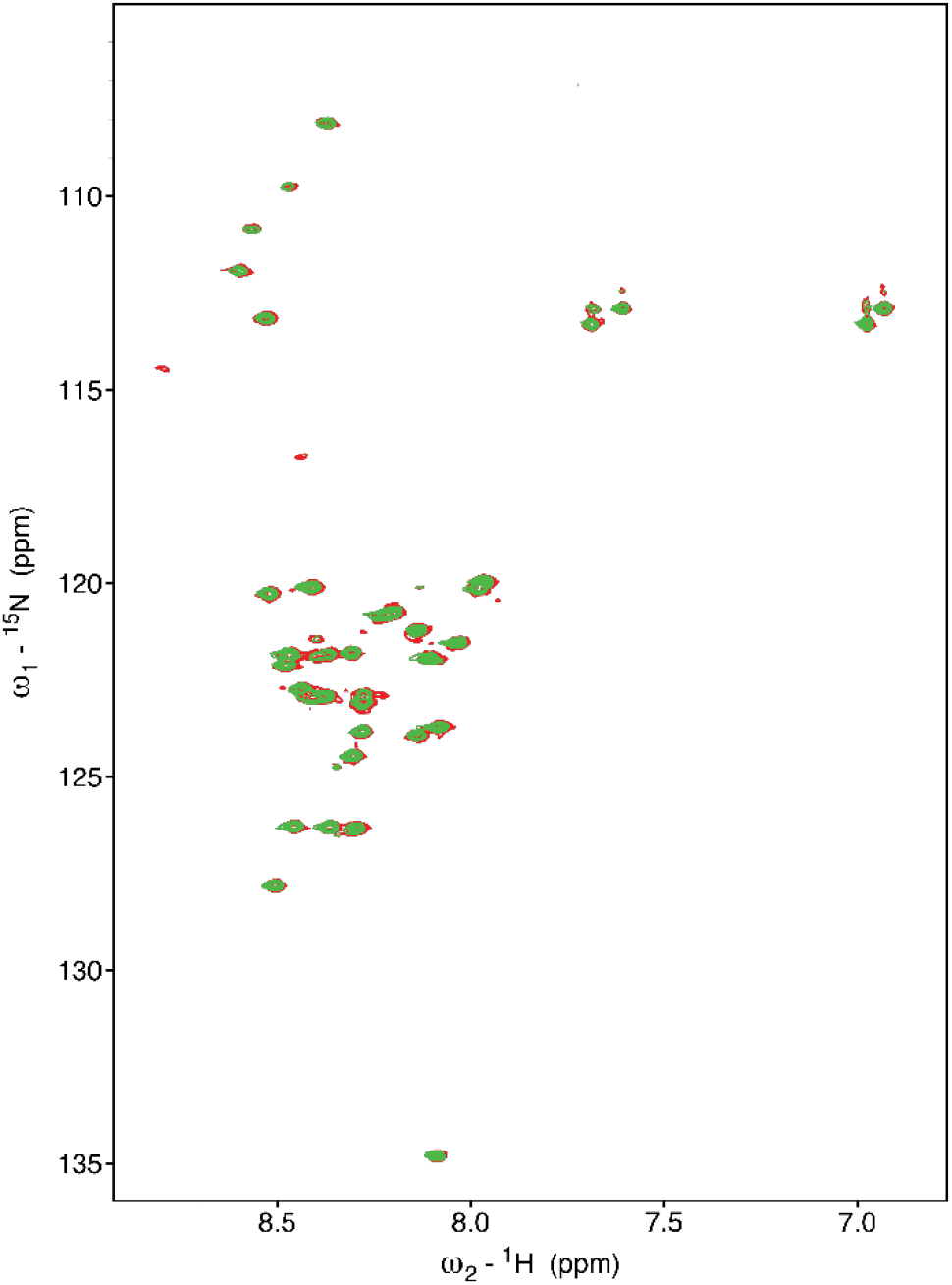
^1^H-^15^N HSQC spectrum of 50 µM Aβ42 at 10 °C. The data was processed using iterative soft thresholding reconstruction method (non-uniformly sampled at 50%). No chemical shift changes were observed in the presence of 200 µM imipramine.

## Conclusion

Our results show that imipramine binds to Aβ42 but not Aβ40 monomers in vitro, setting the stage for follow up in vitro and in vivo studies of its action as a potential competitive ligand in the presence of ApoE.

## Methods

### NMR spectroscopy and sample conditions

NaOH-pretreated Aβ42 (rPeptide) and Aβ40 (Peptide 2.0) were prepared at 20-50 µM (as indicated in figure legends for each type of measurement). Imipramine hydrochloride (Sigma-Aldrich) was prepared at 200 µM. We utilized DPBS buffer containing 0.05% NaN3 (w/v) and 10% D2O (v/v). All NMR measurements were performed at 14.1 T spectrometers (University of Colorado and Denver University) equipped with cryoprobes. STD spectroscopy was performed as described previously^4^. Briefly, NMR saturation was applied on-resonance at 0.65 ppm (corresponding to Aβ resonances), and at -40 ppm as a control (off-resonance). The saturation time illustrated in Fig. 2 was 2 s. A total of 256 scans per measurement were collected and the measurements were performed at 4, 10, and 20 °C. Two-dimensional NOE measurements were collected using the mixing time of 300 ms. 32 scans, 8192 x136 complex points, and States-TPPI acquisition in the indirect dimension were employed.

^1^H-^15^N HSQC experiments were acquired with a nonuniform sampling (NUS) scheme generated by the NUS@HMS scheme generator employing 1024 complex data points in the direct dimension and 50% sampling of the original 200 complex points in the indirect ^15^N dimension. The spectral widths were 13.7 and 35 ppm for the ^1^H and ^15^N dimensions, respectively, with a relaxation delay of 1.3 s and 32 scans. Data reconstruction of the 2D NUS spectra was performed using the hmsIST software.^7^

All 2D NMR data processing and visualization were conducted using NMRpipe/NMRDraw.^8^ Spectra analysis was carried out using the CCPNmr analysis software v2.5.1.^9^ One dimensional STD data was processed in Matlab.

## Acknowledgements

This work is supported by NIH grant RF1AG078965. Some of the measurements utilized the NMR spectrometer at Denver University, supported by National Science Foundation grant DBI-2320158.

## Notes

### Competing Interest Statement

The authors have declared no competing interest.

