## Supporting Information for "Imipramine binds to Amyloid-β (1-42) monomers in vitro, as shown by NMR spectroscopy"

**Supplementary Data: Imipramine binds to Amyloid-β (1-42) monomers in vtiro, as shown by NMR spectroscopy.**

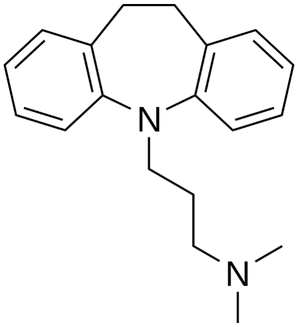


**Figures S1.** Chemical structure of imipramine.

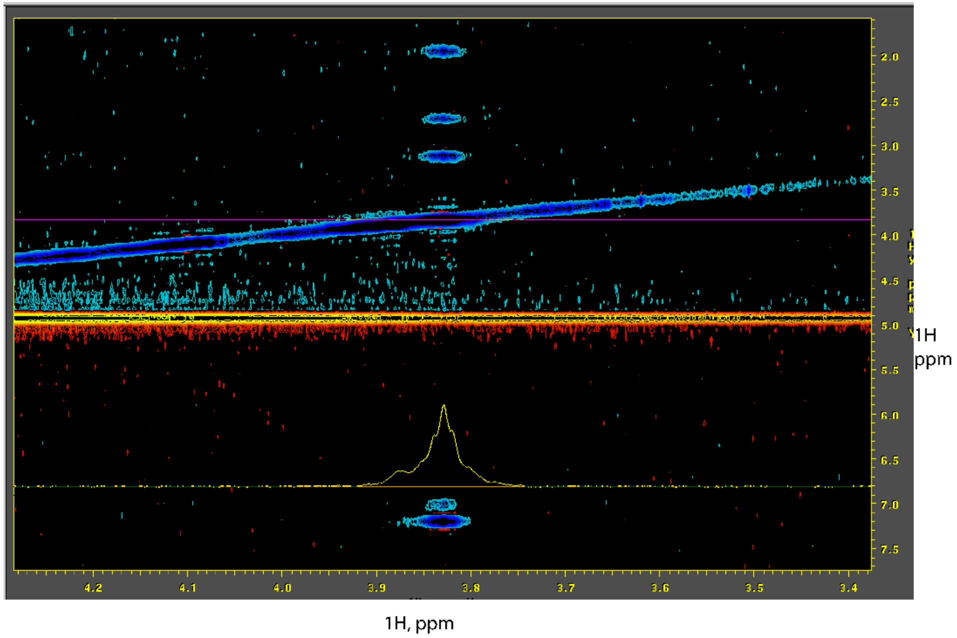


**Figure S2.** Zoom-in region of $^{1}H$-$^{1}H$ NOE of imipramine peaks in the presence of Aβ42, from Figure 1B of the main text. The broadening of the peaks is an additional confirmation of the binding.

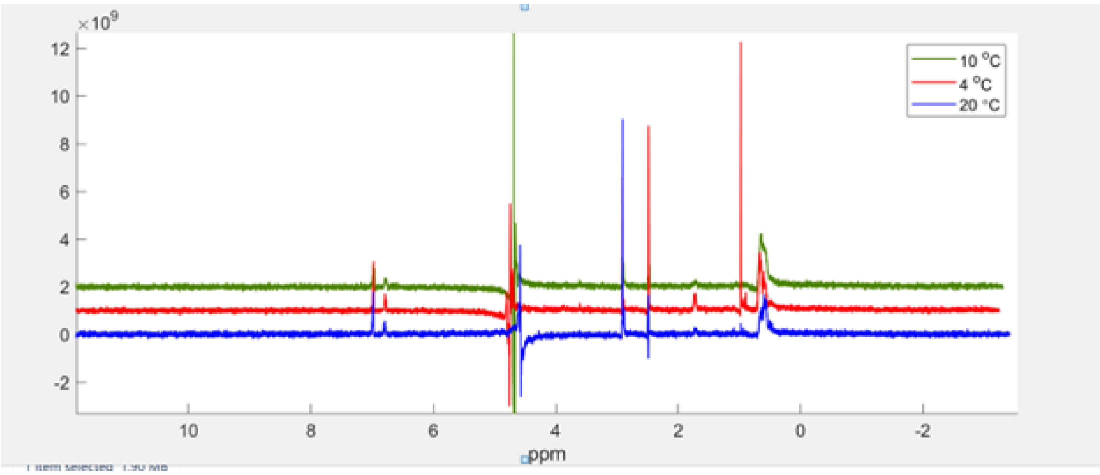


**Figure S3.** NMR measurements of imipramine and Aβ42 binding. a) [1]H NMR STD spectrum for 200 μM imipramine with 20 μM Aβ42 at 4 °C (red), 10 °C (green), and 20 °C (blue).

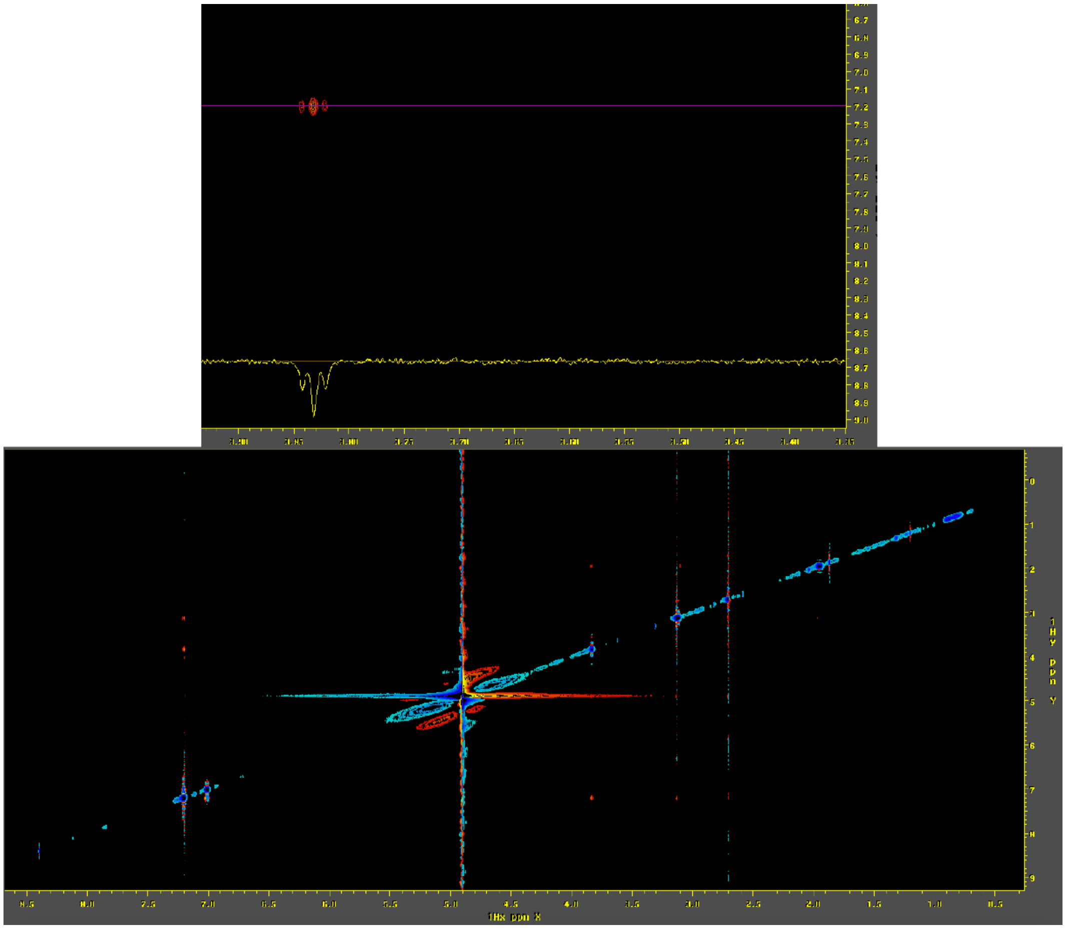

**Figure S4.** $^1$H-$^1$H NOE spectra of imipramine at 200 μM concentration in the presence of 22 μM Aβ40 monomers, at 10°C. The mixing time was 300 ms. The insert on top shows well-resolved imipramine peaks of the opposite sign to the diagonal, excluding binding.
